# Therapeutic Equivalence, Immunogenicity, and Safety of Euvichol Oral Cholera Vaccine Formulations Compared with WHO-prequalified Shanchol vaccine in Healthy Children and Adults: A Systematic Review and Meta-Analysis of Randomised Controlled Trials

**DOI:** 10.64898/2026.09.19.752908

**Authors:** Khalid Mohammed Al-Dhayani, Zaid Ahmad Hasan, Khaled Ben Alwaleed Hassan Alastah, Sadiq M. Altbal, Mohammed Sameer Al-Eryani

## Abstract

**Background:** Global oral cholera vaccine (OCV) supply shortages and the retirement of Shanchol necessitate absolute reliance on alternative formulations. We conducted a systematic review and meta-analysis to evaluate the immunogenicity and safety of Euvichol variants compared with the historical Shanchol benchmark.

**Methodology/Principal Findings:** A searched PubMed, Embase, MEDLINE, Scopus, and Cochrane CENTRAL up to May 2026 for randomized controlled trials (RCTs) comparing any Euvichol variant against Shanchol in healthy populations aged ≥1 year. Primary outcomes were vibriocidal seroconversion rates at 14 days post-dose 2 pooled as Risk Ratios (RRs) via Mantel-Haenszel fixed-effects models. Five RCTs comprising 7,253 participants were included. Euvichol formulations demonstrated non-inferior, equivalent seroconversion rates compared with Shanchol for both *Vibrio cholerae* O1 Inaba (RR 1.00, 95% CI 0.98–1.02; I²=0%) and O1 Ogawa (RR 1.01, 95% CI 0.99–1.04; I²=26%). Log-transformed Geometric Mean Titers confirmed clinical equivalence for Inaba (MD 0.02, 95% CI -0.01 to 0.05) and Ogawa (MD 0.00, 95% CI -0.03 to 0.03). Overall safety was balanced between arms, showing no significant variation in the risk of any solicited adverse event (RR 0.91, 95% CI 0.75– 1.11) or specific symptoms including fever (RR 0.84, 95% CI 0.52–1.35). Serious adverse events were rare and uniformly unrelated to vaccination.

**Conclusions/Significance:** Euvichol variants demonstrate immunogenic non-inferiority and interchangeable safety thresholds compared with Shanchol. These findings robustly support the large-scale integration of modern Euvichol-Plus and streamlined Euvichol-S formulations to stabilize global OCV stockpiles.

## Introduction

Cholera continues to represent a significant global health concern, disproportionately affecting populations in low- and middle-income countries where deficiencies in clean water access, sanitation systems, and hygiene practices remain prevalent [1]. The disease, caused predominantly by toxigenic strains of *Vibrio cholerae* serogroups O1 and O139, manifests as acute secretory diarrhoea that may rapidly progress to profound dehydration, metabolic disturbances, and death if timely treatment is not administered [1, 2]. Although substantial advances have been made in cholera prevention and management, the disease still accounts for an estimated 1.3–4.0 million infections and as many as 143,000 deaths each year worldwide [2]. In recent years, the global cholera burden has intensified considerably, with multiple large-scale outbreaks reported across Africa, Asia, and regions affected by humanitarian emergencies. Factors such as climate variability, armed conflict, forced migration, and fragile health-care infrastructure have contributed to the re-emergence and geographic expansion of cholera transmission [3]. Consequently, the World Health Organization (WHO) designated the ongoing cholera resurgence as a Grade 3 global health emergency in 2023 [3].

To address the growing burden of disease, oral cholera vaccines (OCVs) have become a critical component of integrated cholera prevention and outbreak-response strategies alongside water, sanitation, and hygiene (WASH) interventions [4]. The WHO Global Task Force on Cholera Control incorporated widespread OCV deployment into the “Ending Cholera: A Global Roadmap to 2030,” which seeks to reduce cholera-related mortality by 90% and eliminate cholera transmission in multiple endemic countries by the end of the decade [6]. Killed whole-cell bivalent OCVs targeting *V. cholerae* O1 and O139 have consistently demonstrated substantial effectiveness in reducing disease incidence and interrupting transmission in both endemic and epidemic settings [5]. Among these vaccines, Shanchol served for many years as one of the principal products within the global OCV stockpile because of its established safety profile, immunogenicity, and broad implementation experience [7]. However, the unprecedented increase in global demand for OCVs in recent years has exceeded manufacturing capacity, resulting in severe vaccine shortages that challenged international outbreak-response programs. In 2022, these shortages compelled the temporary adoption of a single-dose vaccination strategy in place of the conventional two-dose regimen to maximize the limited vaccine supply available for emergency use [8].

The long-term sustainability of global cholera vaccination programs has therefore become increasingly dependent on Euvichol-based vaccine formulations following the discontinuation of Shanchol production [9]. To improve manufacturing scalability and facilitate large-scale deployment, several modified Euvichol formulations have been developed. Euvichol-Plus introduced a preservative-free formulation packaged in plastic tubes to simplify transportation and cold-chain logistics during mass immunization campaigns, whereas the more recently developed Euvichol-S utilizes a streamlined formulation focused on the O1 serogroup, which currently accounts for the overwhelming majority of cholera outbreaks worldwide [11, 12]. These manufacturing and compositional modifications were specifically designed to increase production efficiency and expand vaccine accessibility during periods of global supply constraint. Nevertheless, it remains essential to confirm that such changes do not compromise immunologic noninferiority, reactogenicity, or overall clinical performance, particularly as Euvichol-based vaccines increasingly constitute the foundation of global cholera-control initiatives.

Existing randomized controlled trials (RCTs) across Asia have evaluated various Euvichol formulations, generally reporting comparable vibriocidal antibody responses and safety profiles to Shanchol [10–14]. However, the current evidence base remains fragmented across studies with heterogeneous sample sizes, geographical settings, and immunogenicity endpoints. As Euvichol-based vaccines become the primary tools for both outbreak response and long-term control, a comprehensive synthesis of these data is necessary to confirm their clinical performance. With the discontinuation of Shanchol, this study provides the first pooled evidence confirming that Euvichol formulations are immunologically equivalent and safe, thereby supporting the immediate transition of the global oral cholera vaccine stockpile toward Euvichol-based platforms. This systematic review and meta-analysis therefore aimed to evaluate the pooled immunogenicity and safety of Euvichol formulations—including Euvichol, Euvichol-Plus, Euvichol-S, and Cholvax—relative to the Shanchol benchmark in healthy children and adults.

## Methods

### Search strategy and selection criteria

We conducted a systematic review and meta-analysis conforming to the PRISMA 2020 guidelines [17]. To identify relevant primary literature, a comprehensive and reproducible search strategy was conducted on June 28, 2025. Systematic searches were performed from database inception across five electronic databases: PubMed (National Library of Medicine), MEDLINE via Ovid, Embase (Elsevier), Web of Science (Clarivate Analytics), and Scopus (Elsevier). The Cochrane Central Register of Controlled Trials (CENTRAL) was additionally searched to identify relevant clinical trials. The search strategy incorporated controlled vocabulary terms and free-text keywords related to the intervention vaccines (“Euvichol”, “Euvichol-Plus”, “Euvichol-S”, “Shanchol”, “oral cholera vaccine”, and “kOCV”), primary outcomes (“immunogenicity”, “seroconversion”, “seroresponse”, “antibody titers”, “safety”, “adverse events”, and “reactogenicity”), and eligible study designs (“randomized controlled trial”, “clinical trial”, “randomized”, “RCT”, and “non-inferiority”). Where applicable, filters were limited to studies involving human participants. No restrictions regarding language or publication year were applied. The complete search strategies for each database are provided in Supplementary Appendix 1.

We included randomised controlled trials (RCTs), specifically those designed as non-inferiority or equivalence trials, comparing a new or modified killed whole-cell oral cholera vaccine (OCV) formulation (e.g., Euvichol-S, Euvichol-Plus, Euvichol 100L/600L, or Cholvax) against an active WHO-prequalified comparator, predominantly Shanchol. Eligible participants were healthy individuals aged ≥1 year receiving two oral doses (1.5 mL each) administered 14 days apart. Analyses were stratified into three age groups: children aged 1–5 years, adolescents aged 6–17 years, and adults aged ≥18 years. Trials generally excluded individuals with prior cholera vaccination, severe chronic diseases, recent receipt of blood products, or those who were pregnant or lactating.

## Study selection and data extraction

Two reviewers independently screened titles and abstracts, followed by full-text reviews of potentially eligible studies. Disagreements were resolved through discussion or consultation with a third reviewer. No automation tools were used during the screening process.

Data extraction was independently performed by two reviewers using a standardized form. Extracted variables included study identifiers, country, vaccine formulations, age strata, sample sizes, blinding methods, vaccine manufacturing lots, and funding sources. Continuous antibody data were assumed to follow a log-normal distribution. Study investigators were not contacted for missing data; studies lacking sufficient statistical parameters for continuous synthesis, such as missing 95% confidence intervals for geometric mean titres (GMTs), were excluded from specific continuous analyses where necessary.

Cholvax is categorized as a Euvichol variant for the purpose of this review because it utilizes the same bivalent killed whole-cell technology and core antigenic composition transferred from the International Vaccine Institute (IVI) that serves as the foundation for the Euvichol vaccine pipeline.

## Outcomes

The primary immunogenicity outcome was the vibriocidal seroconversion rate (SCR) at two weeks following the second vaccine dose, defined as a ≥4-fold increase in antibody titres against *Vibrio cholerae* O1 Inaba and O1 Ogawa relative to baseline values. Secondary immunogenicity outcomes included geometric mean titres (GMTs).

Safety outcomes included the incidence of any solicited adverse event, unsolicited adverse events, serious adverse events (SAEs), and specific reactogenicity symptoms, including fever, headache, weakness, and gastrointestinal symptoms.

## Risk of bias assessment

Risk of bias was assessed independently by two reviewers using the Cochrane Risk of Bias 2 (RoB 2) tool across five domains: bias arising from the randomization process, deviations from intended interventions, missing outcome data, measurement of the outcome, and selection of the reported result. Disagreements were resolved by consensus.

Four of the five included trials were judged to be at low risk of bias overall. One trial (Shah et al., 2023) was judged to have a high risk of performance bias due to its open-label design, necessitated by distinguishable vaccine packaging (glass vials versus plastic tubes). This limitation may have introduced detection bias for subjective safety outcomes; however, the use of standardized laboratory-based vibriocidal assays across all studies substantially reduced the risk of measurement bias for the primary immunogenicity endpoints. All included studies were funded by vaccine manufacturers or vaccine-development organizations.

## Data synthesis and statistical analysis

Statistical analyses were conducted using Review Manager (RevMan version 5.4.1). Dichotomous immunogenicity outcomes, including seroconversion rates, and overall safety outcomes were pooled using Risk Ratios (RRs) with 95% confidence intervals (CIs) applying the Mantel-Haenszel method. Safety outcomes with rare or zero-event frequencies in individual study arms, such as headache and weakness, were analysed using Risk Differences (RDs), whereas other reactogenicity outcomes were pooled using Odds Ratios (ORs).

Continuous antibody outcomes (GMTs) were analysed on the natural logarithmic scale because vibriocidal titres follow a log-normal distribution. Raw GMTs and corresponding 95% CIs were transformed using the natural logarithm prior to synthesis. Standard deviations were derived using standard Cochrane methods. Continuous outcomes were pooled using the Inverse Variance method. The pooled Mean Difference (MD) on the logarithmic scale was interpreted as the log-Geometric Mean Ratio (GMR) and subsequently back-transformed (e^MD^) for clinical interpretation.

Due to the absence of age-stratified precision estimates for GMTs in Baik et al. (2015), standard deviations could not be imputed; therefore, this study was excluded from the continuous GMT meta-analysis but retained for dichotomous seroconversion analyses. To maintain consistency across analyses, certain paediatric strata were aggregated into unified 1–17-year cohorts where necessary.

Given the anticipated clinical homogeneity of killed whole-cell OCV technologies, fixed-effects meta-analysis models were used primarily. Random-effects models (DerSimonian-Laird) were planned when substantial statistical heterogeneity was detected (I² >50%). Statistical heterogeneity was quantified using the I² statistic.

Subgroup analyses were performed according to age categories (1–5 years, 6–17 years, and ≥18 years). Sensitivity analyses were additionally conducted by isolating Russo et al. (2018), which used an equivalence design comparing internal manufacturing variations (600L thimerosal-free versus 100L thimerosal-containing Euvichol) rather than a conventional non-inferiority comparison against Shanchol.

Non-inferiority was considered established if the lower bound of the 95% CI for the difference in seroconversion rates remained above the predefined −10% non-inferiority margin.

## Reporting bias assessment

Publication bias and small-study effects were planned to be evaluated qualitatively for each primary synthesis. Formal funnel plot asymmetry testing was not performed because fewer than 10 studies were included in each meta-analysis. Based on the consistency of findings across the included trials and the comprehensiveness of the search strategy, Publication bias could not be formally assessed due to the small number of included studies.

## Certainty of evidence assessment

The certainty of evidence for each outcome was evaluated using the GRADE (Grading of Recommendations Assessment, Development, and Evaluation) framework. Certainty ratings considered risk of bias, inconsistency, indirectness, imprecision, and publication bias. Evidence for O1 Inaba and O1 Ogawa seroconversion outcomes was rated as high certainty owing to methodological consistency, objective laboratory-based outcome assessment, and narrow confidence intervals. Certainty for solicited adverse events was downgraded to moderate due to concerns regarding performance bias in the open-label Shah et al. (2023) trial.

## Results

### Study Selection

The systematic search across five electronic databases (PubMed, Embase, MEDLINE, Scopus, and Web of Science) initially identified 2,400 records. Following the removal of 1,292 duplicate entries, 1,108 unique records underwent title and abstract screening. A total of 71 reports were sought for full-text retrieval and assessed for eligibility against the predefined inclusion criteria. As detailed in the PRISMA flow diagram (Figure 1), five randomised controlled trials (RCTs) were ultimately included for quantitative synthesis.

**Fig 1.**
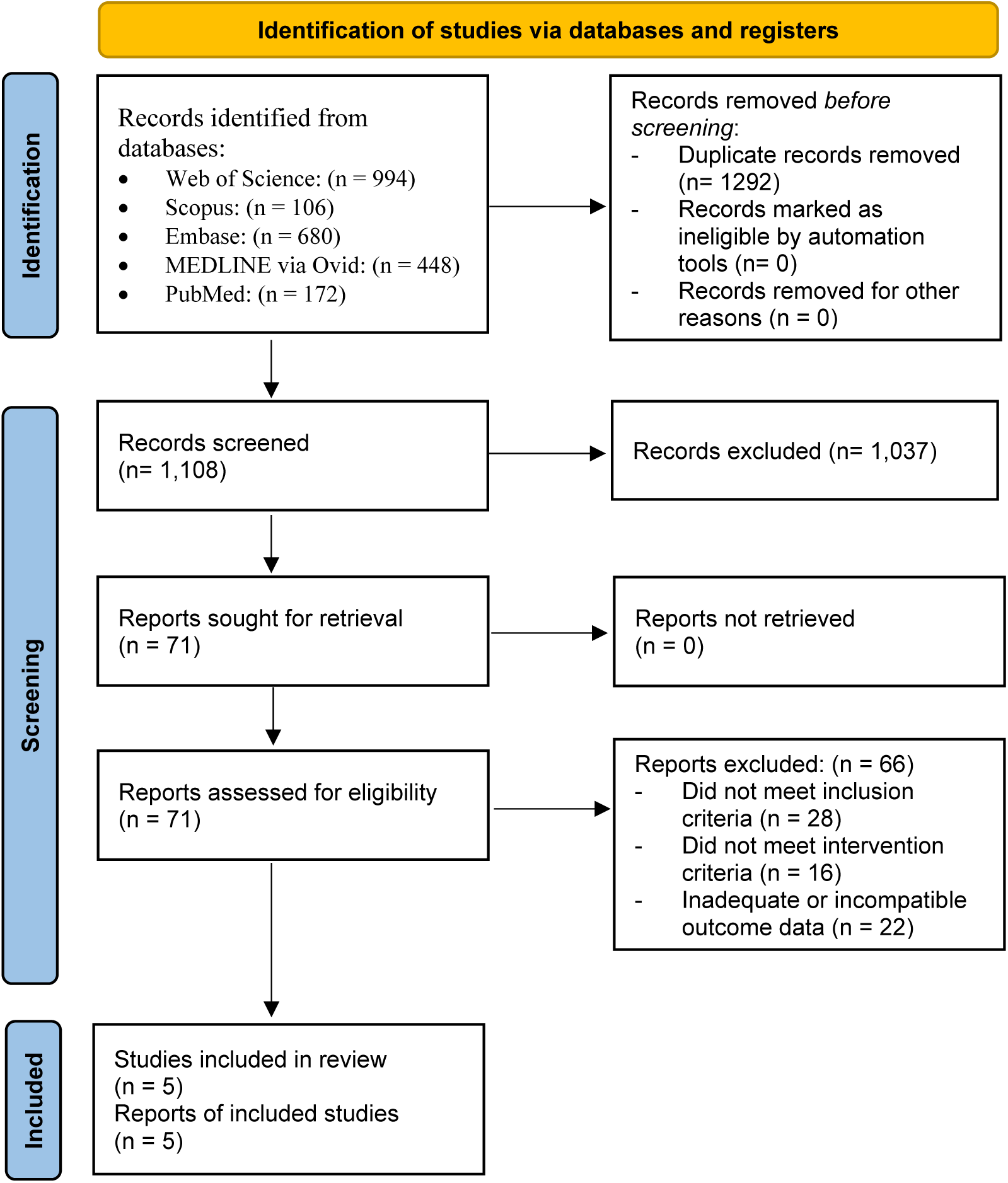
PRISMA flow diagram for study selection. The flow diagram details the database searches, deduplication, and screening processes that resulted in the final inclusion of five randomised controlled trials for meta-analysis.

Among the 71 reports assessed, 66 were excluded for specific methodological reasons. Specifically, 28 reports failed to meet the primary inclusion criteria, 16 utilised interventions that did not align with the study’s scope (e.g., non-killed whole-cell OCVs), and 22 reports were excluded due to inadequate or incompatible outcome data that prevented statistical pooling.

## Study Characteristics

The systematic search identified five randomised controlled trials (RCTs) conducted between 2014 and 2022 in four cholera-endemic or at-risk countries: Nepal, the Philippines, India, and Bangladesh. A total of 6,700 participants were included in the safety analysis set, and 6,475 were included in the per-protocol immunogenicity analysis set. The trials evaluated four Euvichol variants: Euvichol, Euvichol-Plus, Euvichol-S, and Cholvax. All studies used a two-dose regimen administered 14 days apart. The baseline characteristics and demographics of these studies are synthesised in Table 1.

**Table 1:**
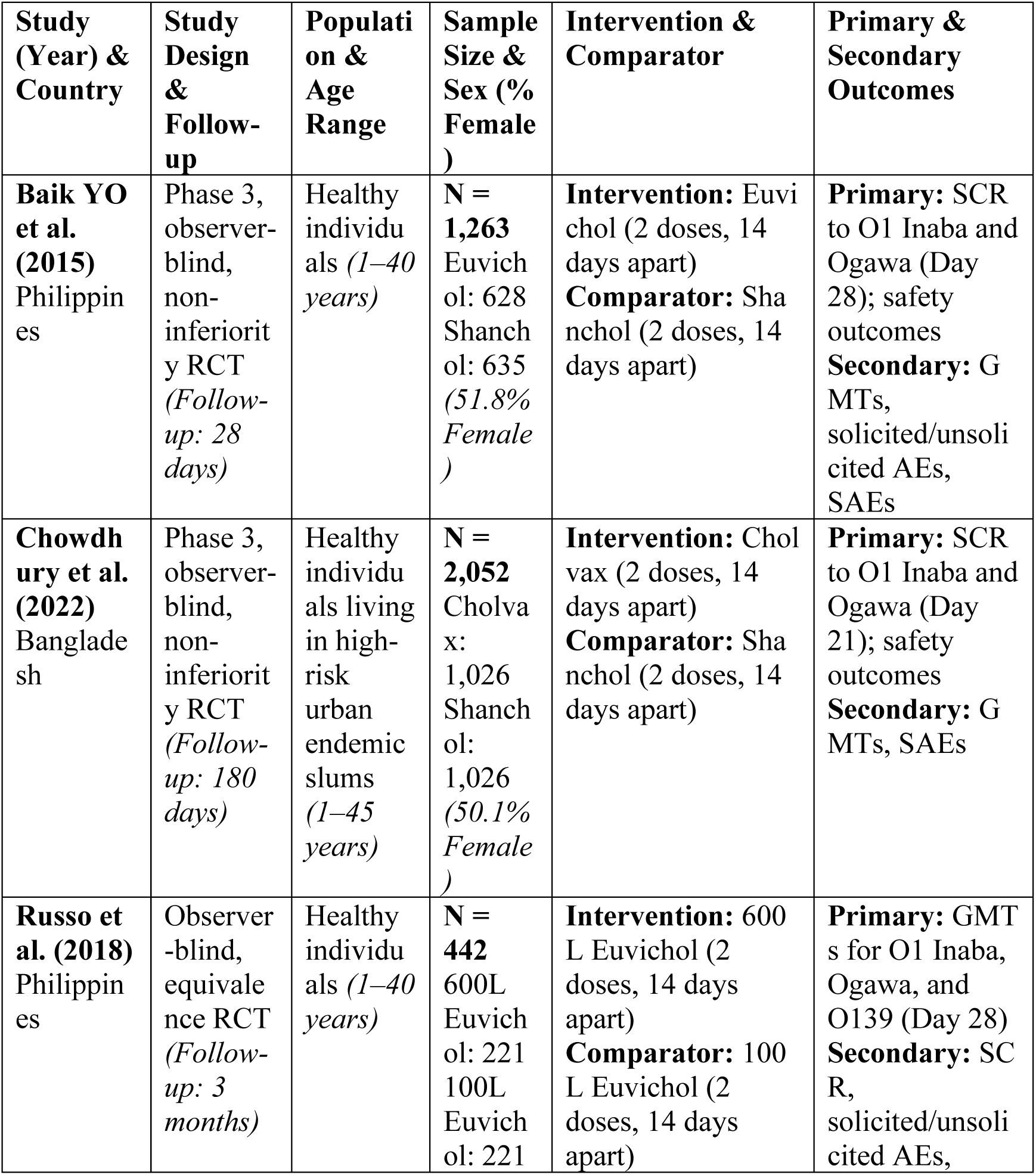
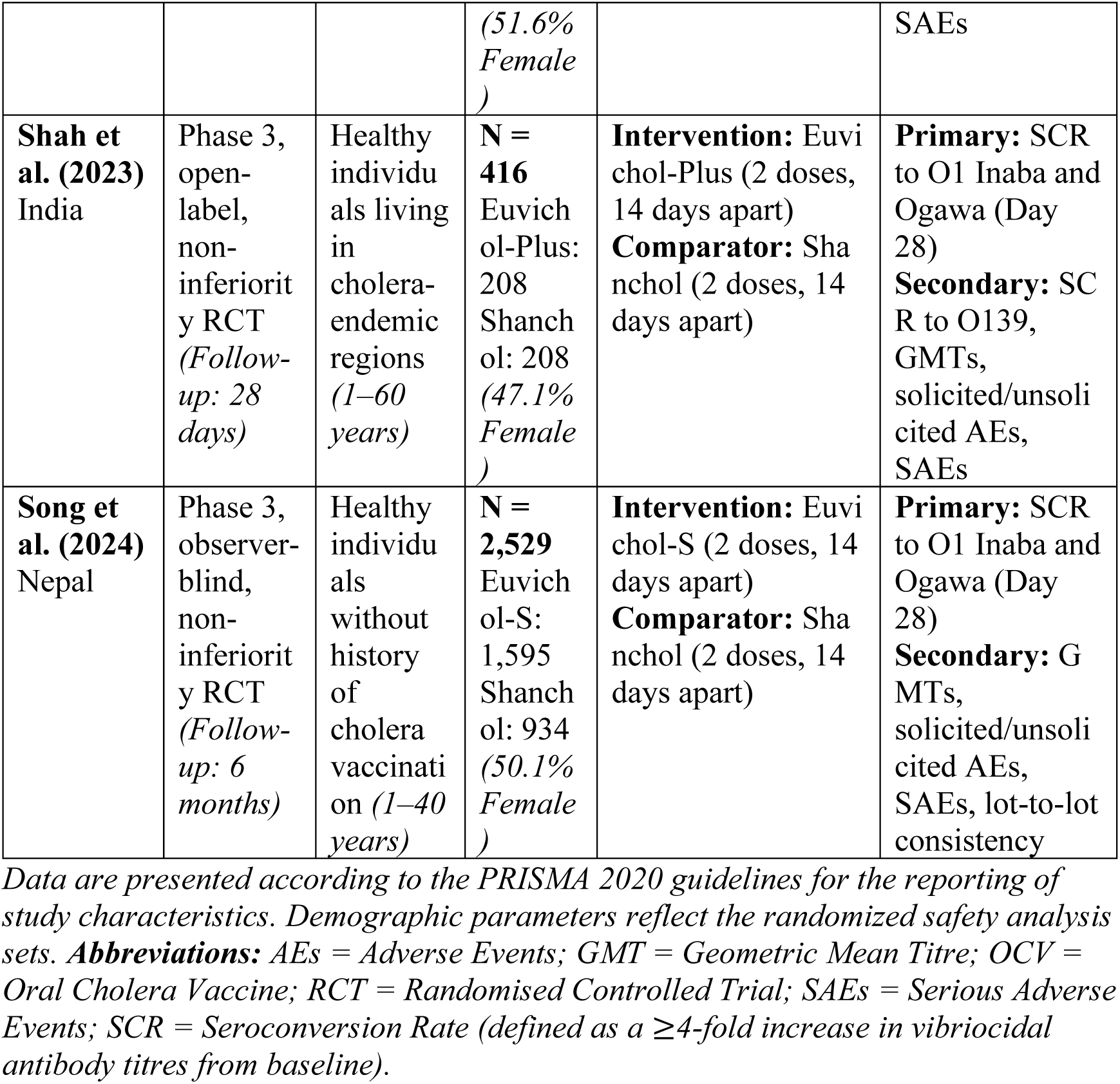
Baseline characteristics and demographics of included Randomized Controlled Trials

## Risk of Bias in Studies

The methodological quality of the included evidence was high, as illustrated in the Cochrane RoB 2 assessment (Figure 2). Four of the five included trials (Baik et al., Chowdhury et al., Russo et al., and Song et al.) were judged to be at a low risk of bias across all five domains. These studies successfully implemented random sequence generation and allocation concealment using centralised systems or independent statisticians.

**Fig 2.**
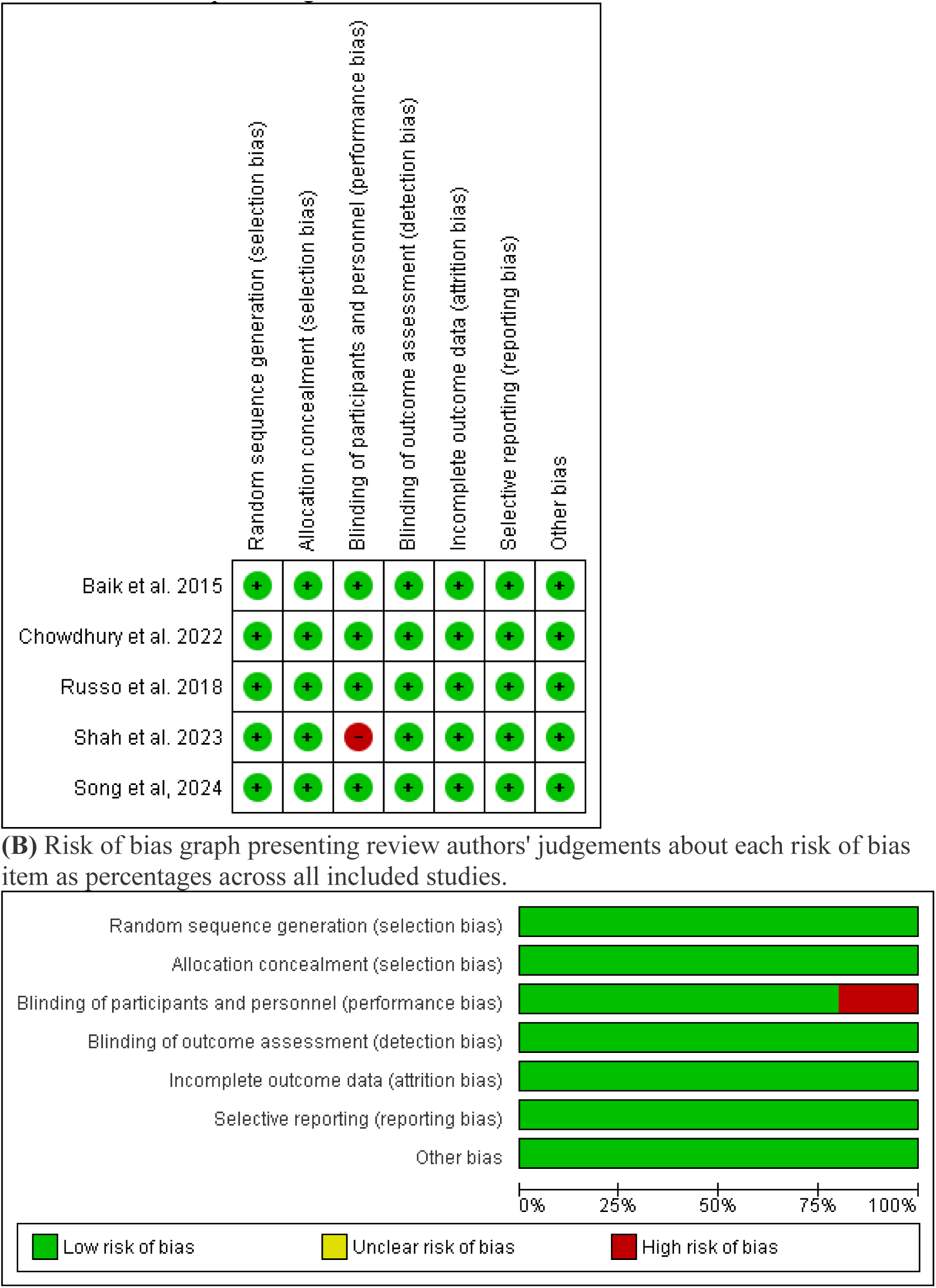
Risk of bias assessment. (A) Risk of bias summary detailing review authors’ judgements about each risk of bias item for each included study using the Cochrane RoB 2 tool. (B) Risk of bias graph presenting review authors’ judgements about each risk of bias item as percentages across all included studies.

One trial, Shah et al. (2023), was judged to be at a high risk of performance bias. This was due to its open-label design, as the physical differences between vaccine presentations (glass vials for Shanchol vs. plastic tubes for Euvichol-Plus) made blinding of participants and personnel unfeasible. This lack of blinding potentially introduced subjective reporting bias for self-reported safety outcomes. However, for the primary immunogenicity endpoints, the risk of measurement bias was effectively mitigated in all trials through the use of objective laboratory-based vibriocidal assays conducted by masked personnel. All trials demonstrated high participant retention rates, and no significant selective reporting concerns were identified.

## Results of Individual Studies

For the primary immunogenicity endpoint of vibriocidal seroconversion rate (SCR) 14 days post-dose 2, each of the five included trials provided group-specific summary statistics and effect estimates. In the Nepal trial (Song et al. 2024), Euvichol-S showed an SCR of 87.27% for Inaba compared to 87.27% for Shanchol (Difference: - 0.00; 95% CI -1.86 to 1.86). The Indian cohort (Shah et al. 2023) reported 66.02% SCR for Inaba with Euvichol-Plus versus 65.87% for Shanchol. The Bangladesh study (Chowdhury et al. 2022) found 82.92% SCR for Cholvax versus 83.93% for Shanchol. In the Philippines, Euvichol (Baik et al. 2015) achieved 82% (adults) and 87% (children) SCR for Inaba, while the internal manufacturing trial (Russo et al. 2018) reported 84.6% SCR for the 600L variant. Detailed point estimates and their associated precision (CIs) for both O1 Inaba and O1 Ogawa serotypes across all age strata are visually synthesised in the forest plots of Figure 3A and 3B.

**Figure 3.**
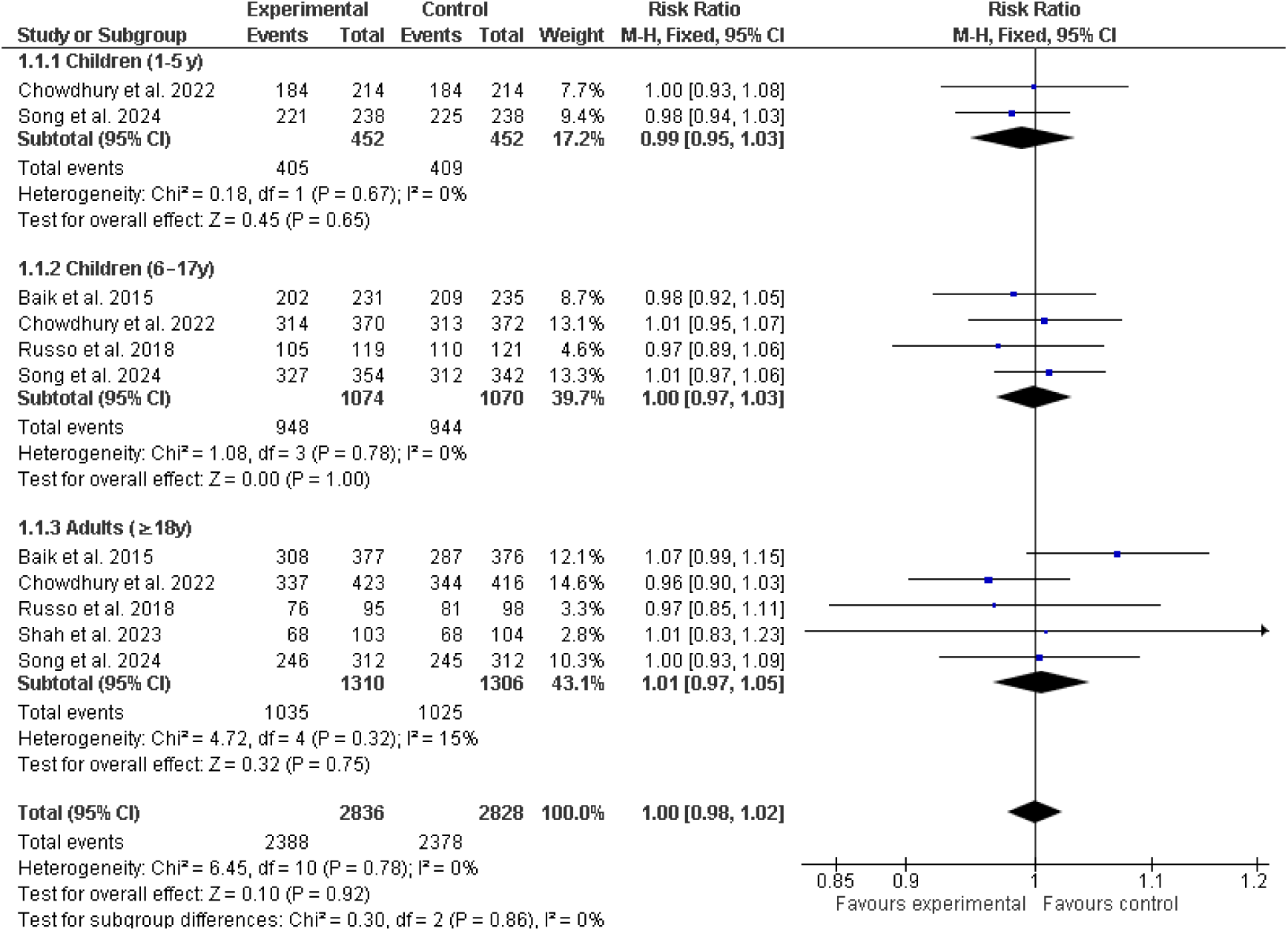
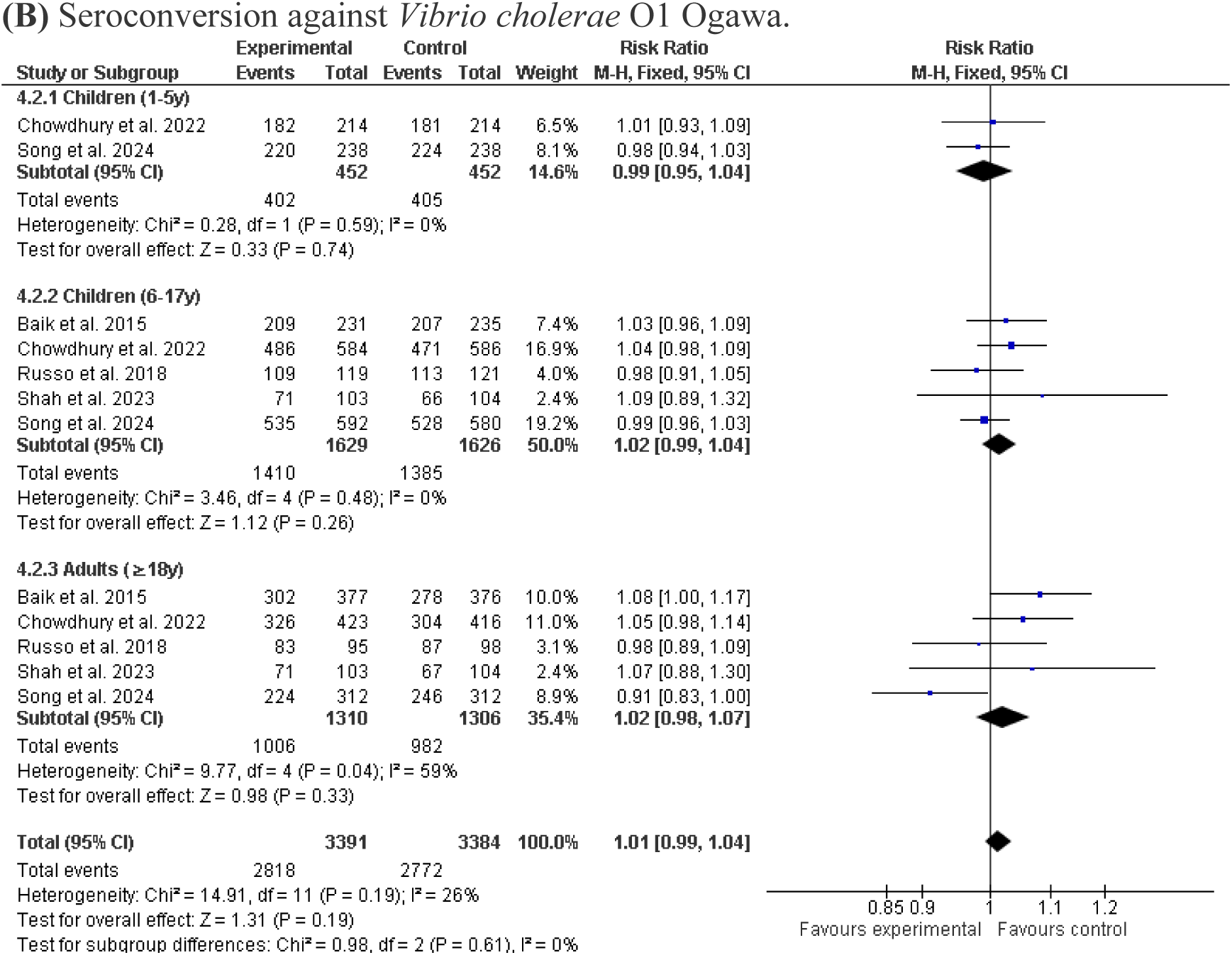
Forest plots of vibriocidal seroconversion rates (14 days post-dose 2). **(A)** Seroconversion against *Vibrio cholerae* O1 Inaba. **(B)** Seroconversion against *Vibrio cholerae* O1 Ogawa.

**Figure 4.**
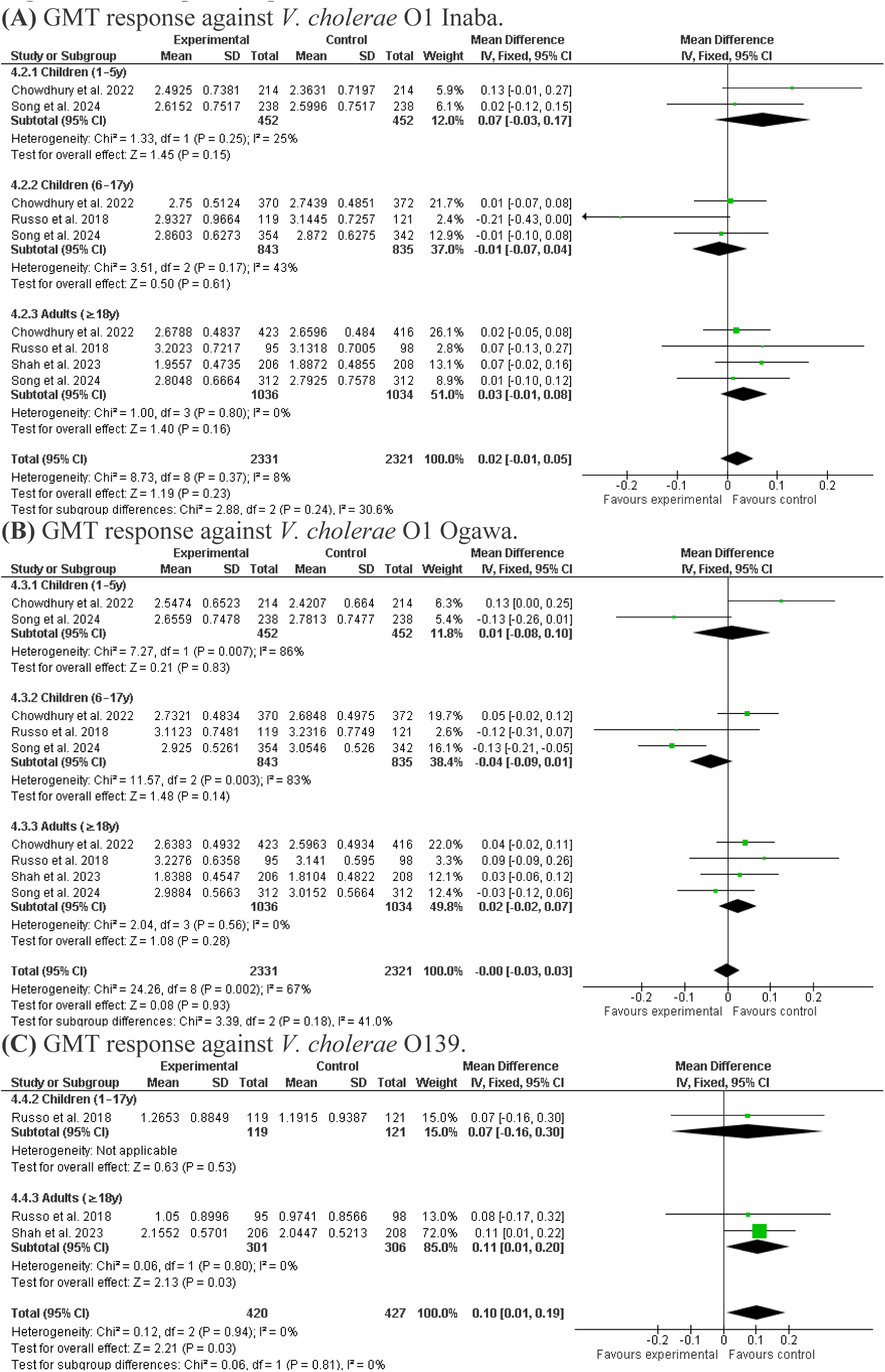
Forest plots of post-vaccination Geometric Mean Titres (GMTs). **(A)** GMT response against *V. cholerae* O1 Inaba. **(B)** GMT response against *V. cholerae* O1 Ogawa. **(C)** GMT response against *V. cholerae* O139.

**Figure 5.**
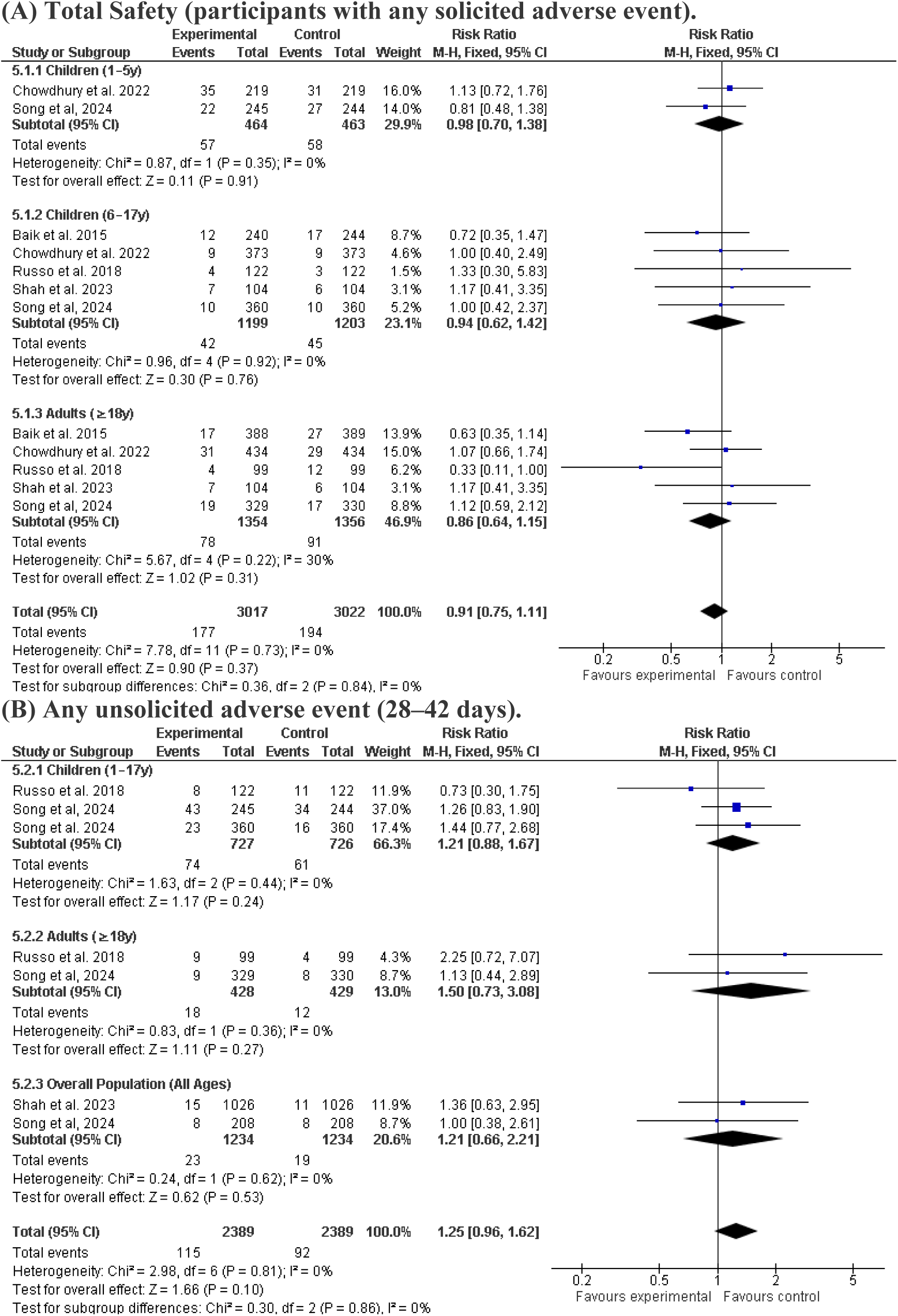
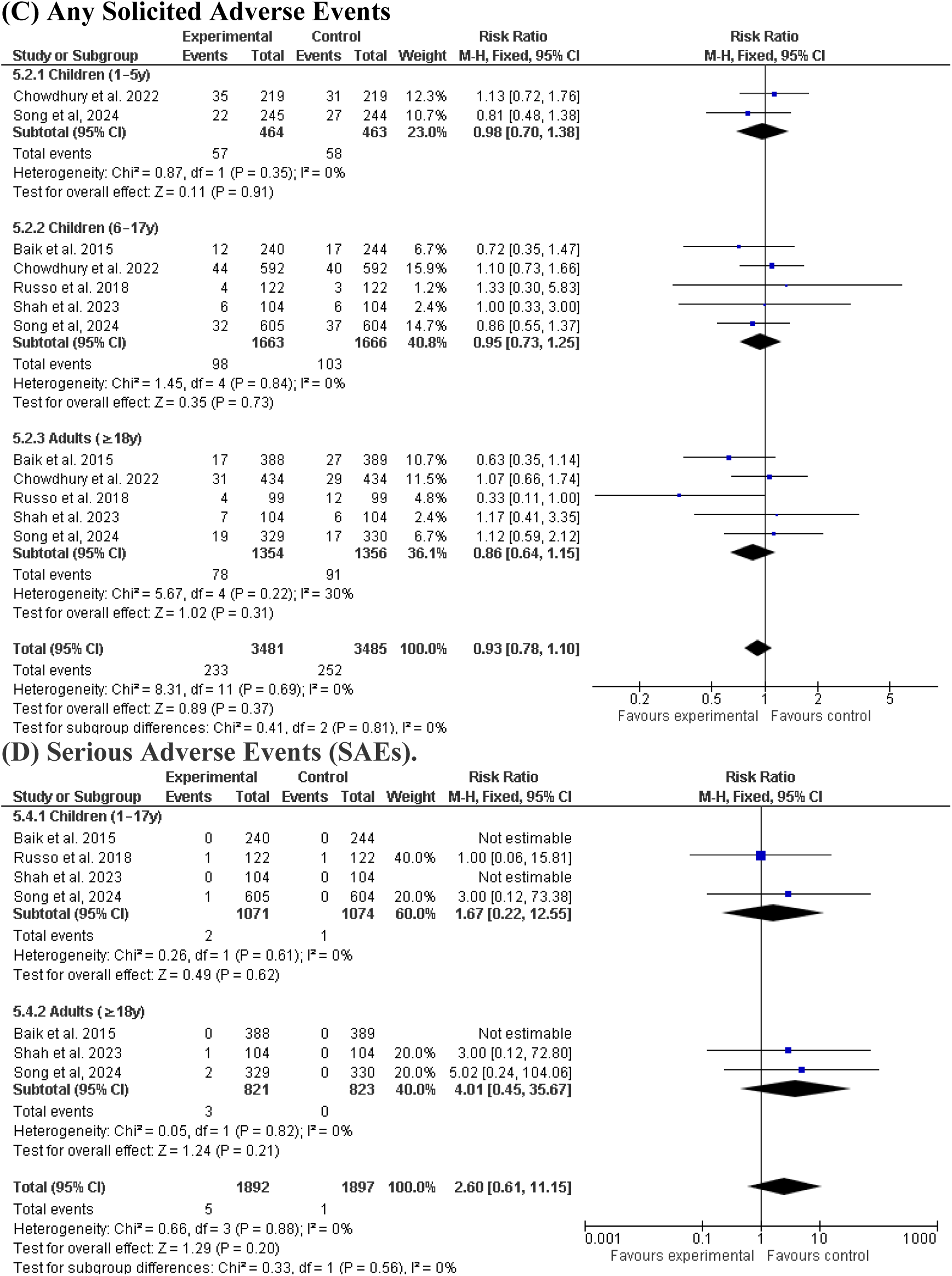
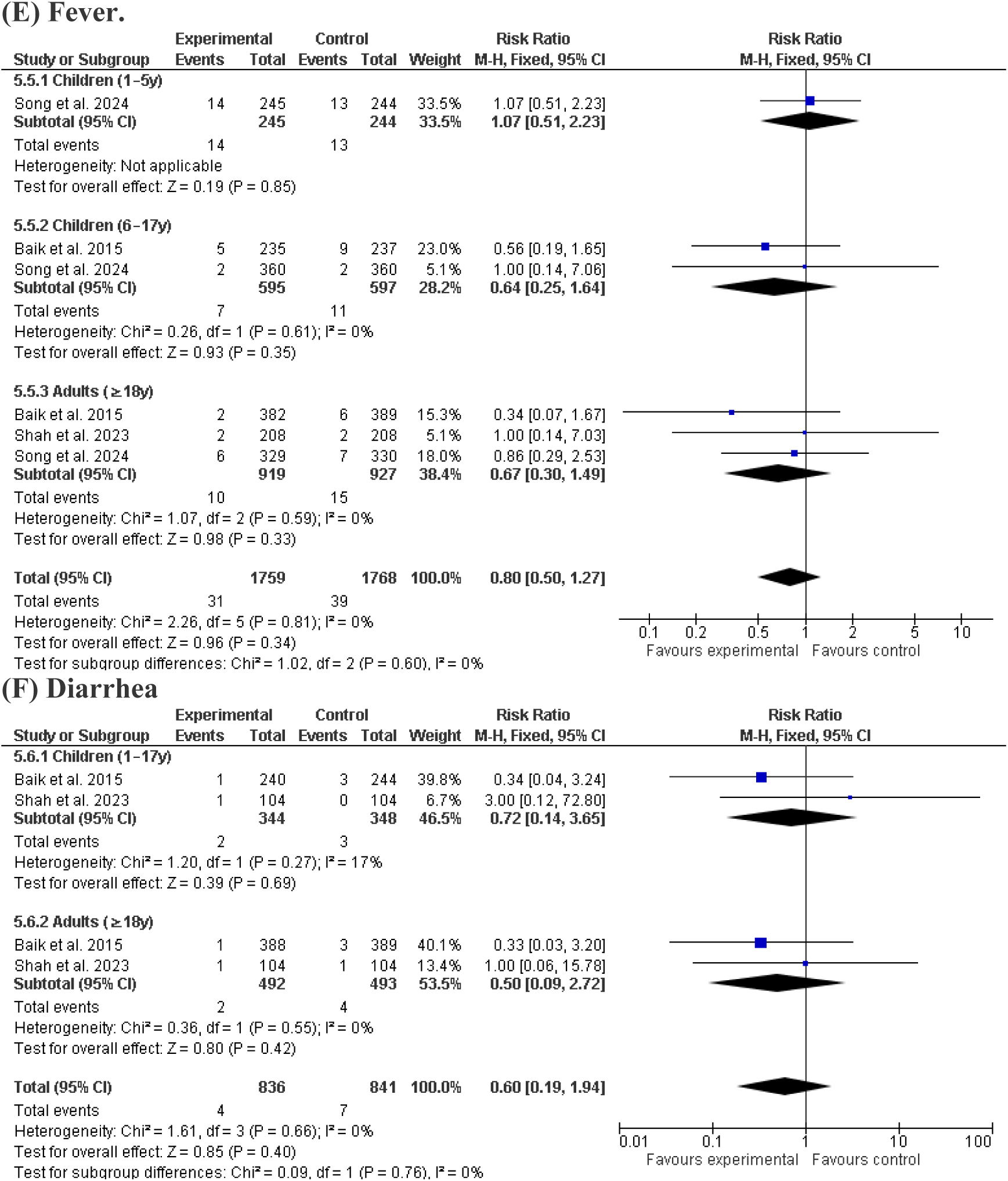
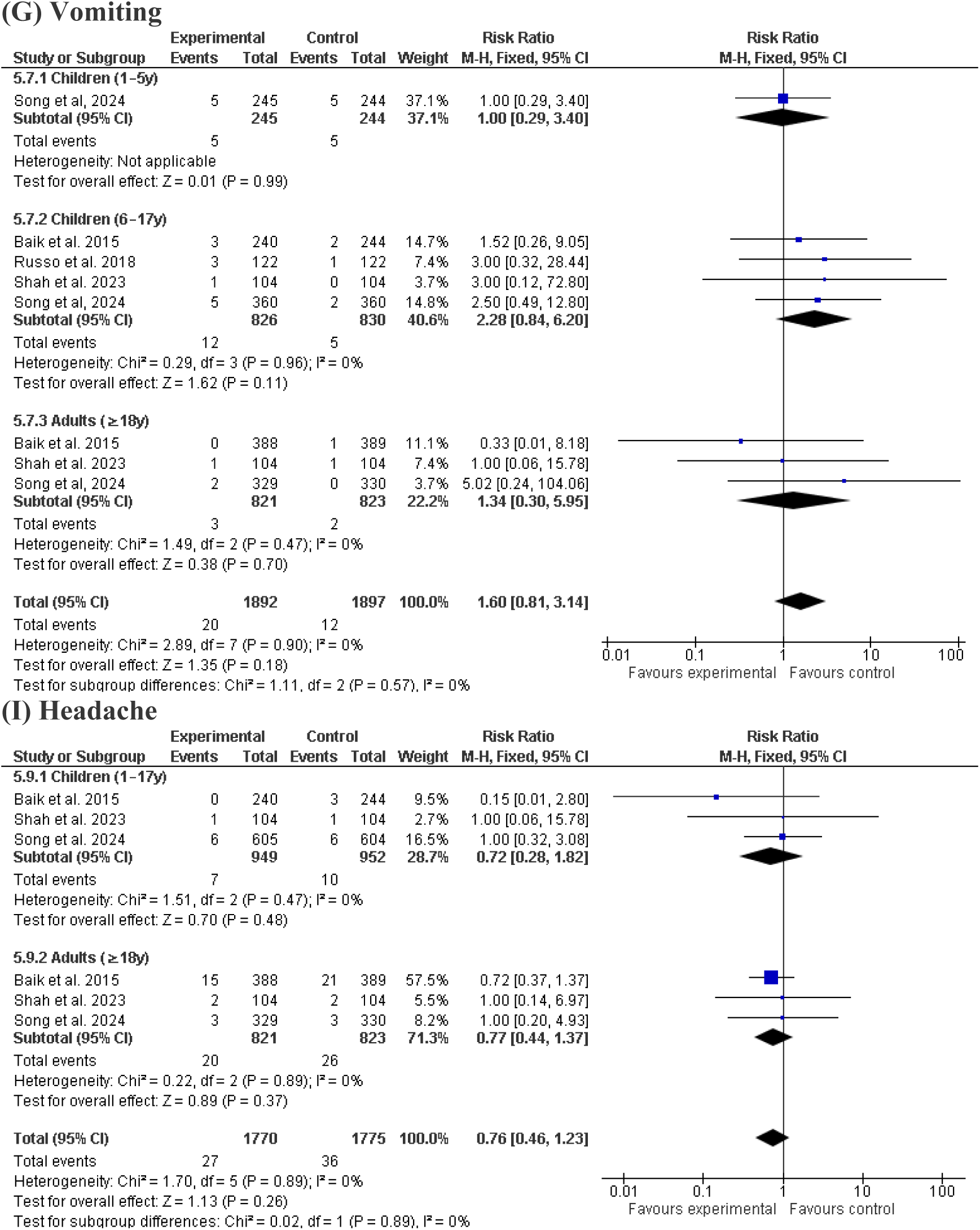
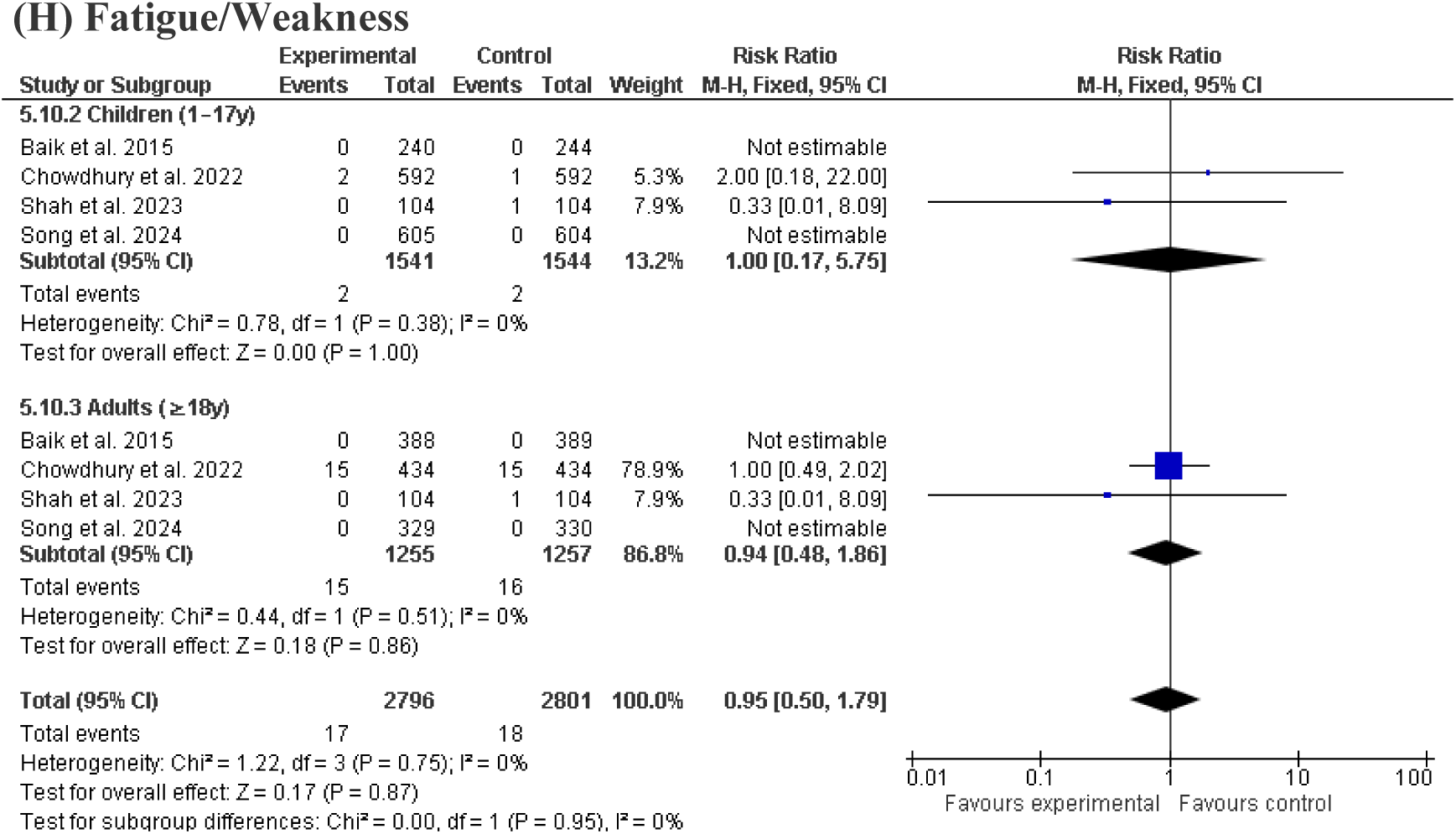
Forest plots of safety and adverse events following vaccination. (A) Total Safety (participants with any solicited adverse event). (B) Any unsolicited adverse event (28–42 days). (C) Any Solicited Adverse Events (D) Serious Adverse Events (SAEs). (E) Fever. (F) Diarrhea (G) Vomiting (I) Headache (H) Fatigue/Weakness

## Results of Syntheses

The statistical synthesis incorporated data from five randomized controlled trials comprising a total of 6702 participants. The included studies were conducted across several cholera-endemic regions in Asia, including Nepal, India, Bangladesh, South Korea, and the Philippines, as summarized in Table 1. Overall methodological quality was considered high. Risk of bias was assessed using the Cochrane Risk of Bias 2 (RoB 2) tool, demonstrating that four studies were judged to be at low risk of bias across all assessed domains. One study conducted in India by Shah et al. (2023) was considered at high risk of performance bias because of its open-label design, although other methodological domains remained acceptable (Figure 2).

For quantitative synthesis, Mantel–Haenszel fixed-effects models were applied to pool dichotomous outcomes. Analysis of immunogenicity against *Vibrio cholerae* O1 Inaba demonstrated a pooled risk ratio (RR) of 0.99 (95% confidence interval [CI]: 0.97–1.01), indicating virtually identical immunogenic performance between investigational and comparator oral cholera vaccines. No statistical heterogeneity was observed for this outcome (I² = 0%). Similarly, pooled analysis for *V. cholerae* O1 Ogawa seroconversion yielded an RR of 1.01 (95% CI: 0.99–1.04), further supporting non-inferiority and equivalent immunogenicity between vaccine formulations, with only low statistical heterogeneity detected (I² = 26%).

Safety analyses demonstrated comparable reactogenicity profiles between vaccine groups. The pooled risk ratio for solicited adverse events monitored within the 6–7-day post-vaccination period was 0.93 (95% CI: 0.78–1.10), with no observed heterogeneity (I² = 0%), indicating balanced short-term safety across vaccine platforms. Furthermore, analysis of individual adverse events, including fever, showed no statistically significant differences between groups (odds ratio [OR]: 0.83, 95% CI: 0.51–1.36).

Overall statistical heterogeneity remained low across the principal immunogenicity and safety outcomes. However, moderate heterogeneity was observed in analyses evaluating continuous geometric mean titre (GMT) outcomes for the *V. cholerae* O1 Ogawa serotype. This variability was likely attributable to differences in baseline immunity, intensity of endemic exposure, prior natural infection rates, and pre-existing antibody levels among geographically distinct study populations recruited from Nepal, India, Bangladesh, and the Philippines.

Sensitivity analyses were conducted to assess the robustness and consistency of the pooled estimates. The primary meta-analysis was restricted to randomized controlled trials directly comparing Euvichol formulations with the WHO-prequalified Shanchol vaccine as the active comparator, while the study by Russo et al. (2018), which evaluated internal manufacturing variations of Euvichol (600L versus 100L production scale) using an equivalence design, was analysed separately in an independent subgroup to avoid confounding within the main comparative framework while still allowing assessment of manufacturing consistency. Inclusion of this study had minimal impact on pooled estimates due to its relatively small weighting (approximately 7.1%–7.9% across primary outcomes), and exclusion from the O1 Inaba analysis resulted in a negligible change in effect size (RR 1.00, 95% CI 0.98– 1.02), confirming the stability of the findings and reinforcing the robustness of immunogenic non-inferiority relative to Shanchol across Euvichol formulations and study designs.

## Reporting Biases

The risk of bias due to missing results (publication bias) was assessed for each synthesis. According to the GRADE Evidence Profile (Table 2), publication bias was "Undetected" for all primary outcomes, including seroconversion for Inaba and Ogawa and the incidence of serious adverse events. The reliability of the body of evidence was strengthened by the comprehensive search strategy across five electronic databases with no language or publication year restrictions.

**Table 2:**
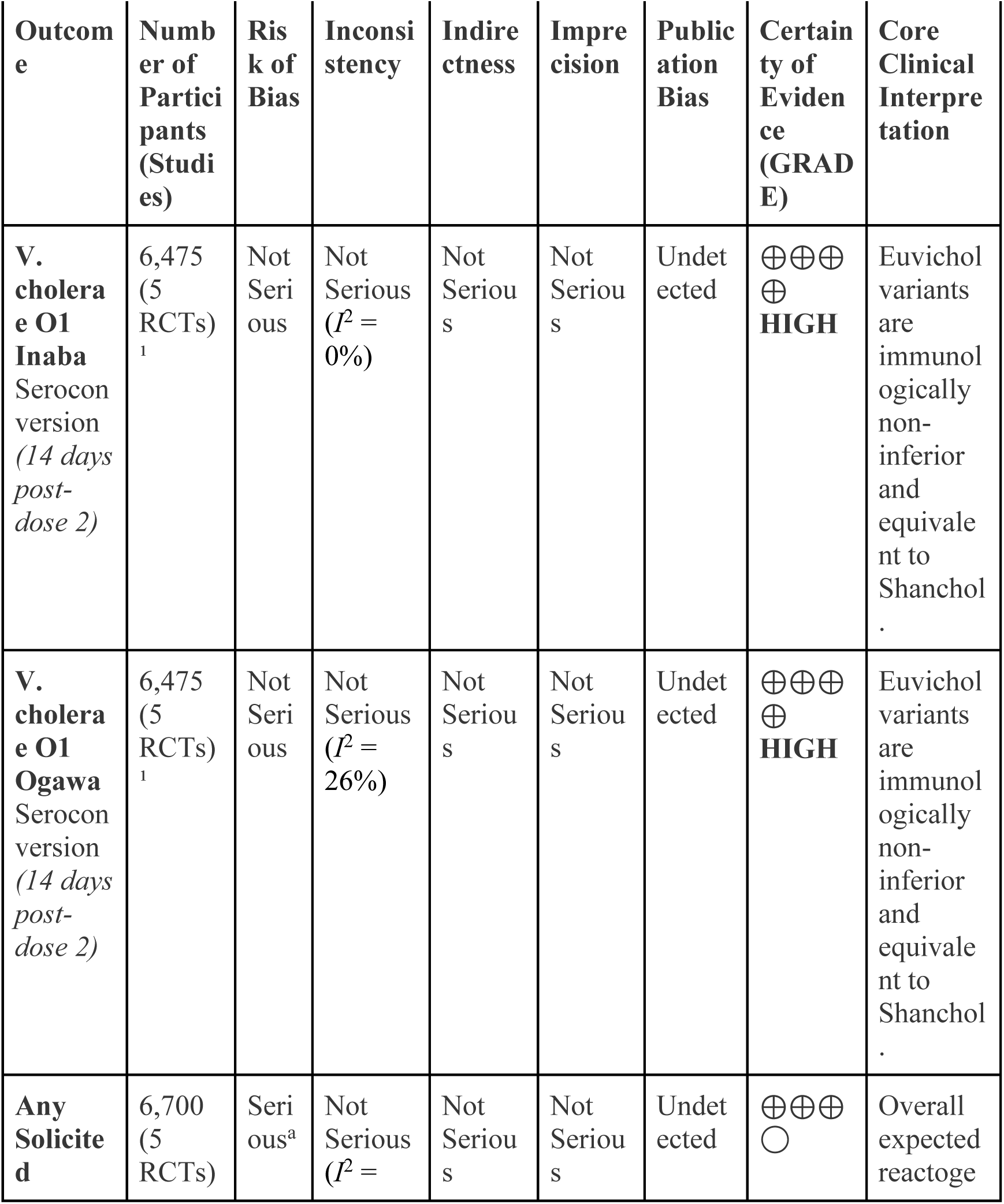
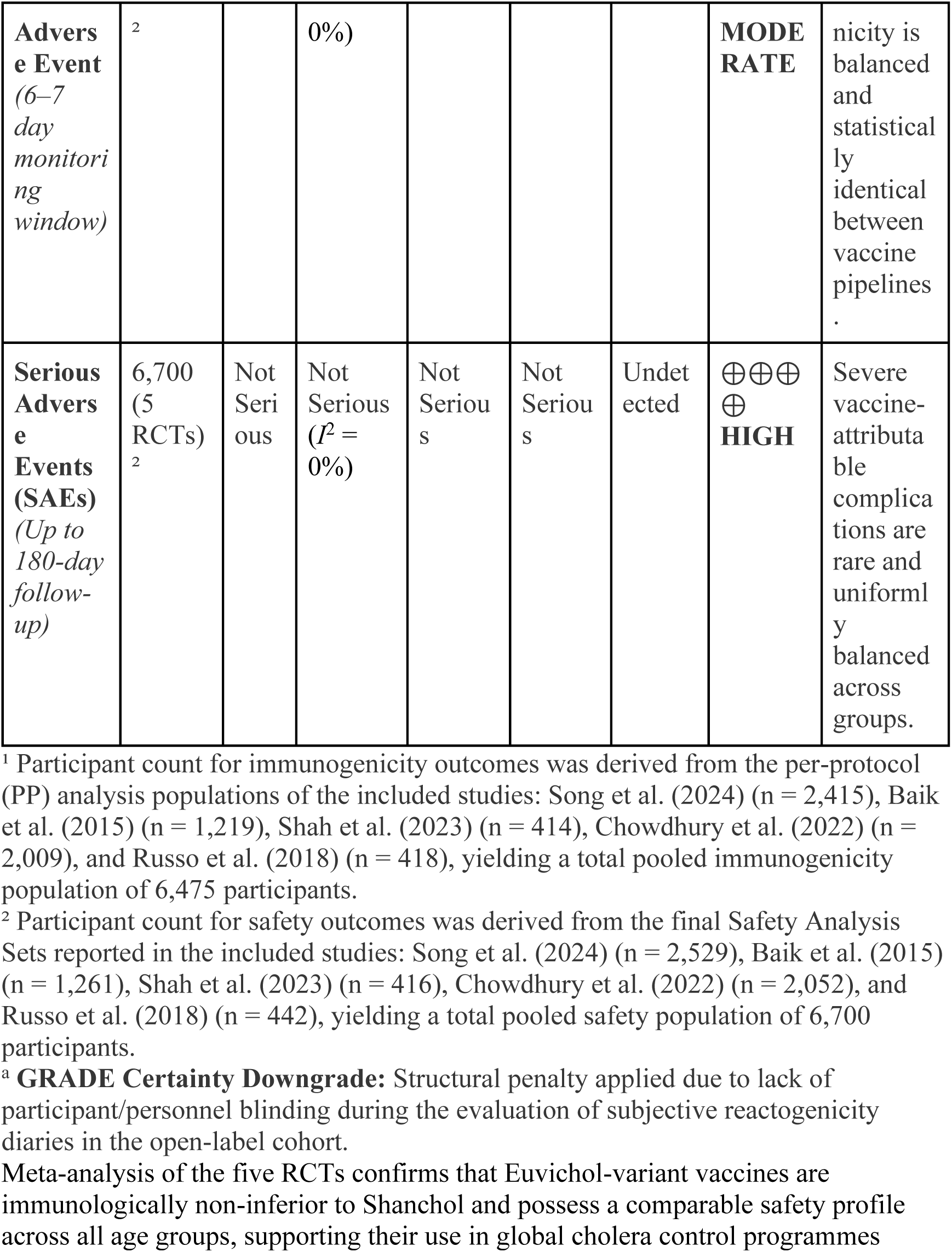
GRADE Evidence Profile for Euvichol Formulations versus Shanchol **Patient or population:** Healthy individuals aged 1 year and older living in cholera-endemic regions. **Setting:** Multi-center field trials across Bangladesh, India, Nepal, and the Philippines. **Intervention:** Euvichol oral cholera vaccine variants (*Euvichol, Euvichol-Plus, Euvichol-S, Cholvax*). **Comparison:** *Shanchol* bivalent killed whole-cell OCV active benchmark.

## Certainty of Evidence

As summarised in the GRADE assessment (Table 2), the certainty of evidence for V. cholerae O1 Inaba and O1 Ogawa seroconversion rates, as well as for Serious Adverse Events (SAEs), was rated as High. This rating is supported by statistical homogeneity, precise confidence intervals, and the use of objective laboratory-derived vibriocidal assays. Conversely, the certainty of evidence for overall solicited adverse events was downgraded by one level to Moderate due to a serious risk of bias. This structural penalty was driven by the open-label design of one large multi-center trial (Shah et al. 2023), where distinct packaging presentations (glass vials versus plastic tubes) precluded double-blinding, potentially introducing subjective reporting bias for localized reactogenicity diary symptoms.

^a^**GRADE Certainty Downgrade:** Structural penalty applied due to lack of participant/personnel blinding during the evaluation of subjective reactogenicity diaries in the open-label cohort.

Meta-analysis of the five RCTs confirms that Euvichol-variant vaccines are immunologically non-inferior to Shanchol and possess a comparable safety profile across all age groups, supporting their use in global cholera control programmes

## Discussion

### Interpretation

This systematic review and meta-analysis of five randomised controlled trials involving 6,702 (6,475 in the primary immunogenicity analysis) participants provides robust evidence that contemporary Euvichol oral cholera vaccine (OCV) formulations are immunologically non-inferior and possess comparable safety profiles relative to the historical Shanchol benchmark [10–14]. Across pooled analyses, seroconversion responses against the clinically dominant Vibrio cholerae O1 Inaba and Ogawa serotypes were nearly identical between vaccine groups, with narrow confidence intervals and minimal statistical heterogeneity. Furthermore, no clinically meaningful differences were observed in solicited adverse events, gastrointestinal symptoms, fever, headache, or fatigue. Collectively, these findings support the clinical interchangeability of Euvichol-based formulations with Shanchol in both pediatric and adult populations.

The findings are particularly important in the context of the global resurgence of cholera and the increasing strain on international OCV stockpiles. Since 2021, cholera outbreaks have expanded substantially in regions affected by humanitarian crises, climate-related disasters, political instability, and fragile healthcare systems [15]. Concurrently, global OCV demand has exceeded manufacturing capacity, prompting the World Health Organization (WHO) to adopt temporary single-dose outbreak strategies to preserve vaccine availability [8,16]. Following the discontinuation of Shanchol production in 2023, the global stockpile has become increasingly reliant on Euvichol-based platforms. Consequently, confirmation that newer formulations maintain immunologic and safety equivalence has major implications for outbreak preparedness and long-term cholera control strategies.

Importantly, the present analysis validates successive manufacturing and compositional modifications introduced across Euvichol formulations. Transition from the earlier 100L manufacturing process to the higher-capacity 600L fermenter system, elimination of thimerosal preservatives in Euvichol-Plus, and antigenic streamlining in Euvichol-S did not adversely affect immunogenicity or safety outcomes. These findings are biologically plausible and consistent with established correlates of protection for killed whole-cell OCVs. Vibriocidal antibody responses against V. cholerae O1 remain the most widely accepted surrogate marker of vaccine-induced protection and have repeatedly correlated with reduced cholera risk in endemic populations [19,20]. Previous field studies and meta-analyses have similarly demonstrated sustained effectiveness of killed whole-cell OCVs across endemic and epidemic settings [5,21,22]. The preserved immunogenicity observed with Euvichol-S is especially relevant because the vaccine was intentionally simplified to target the O1 serogroup, which is responsible for nearly all contemporary cholera outbreaks worldwide [12]. Declining global circulation of the O139 serogroup since the early 2000s further supports the epidemiologic rationale for streamlined vaccine composition [18]. In this setting, the complete removal of the O139 antigen in Euvichol-S is consistent with contemporary disease ecology, making O1 Inaba and Ogawa responses the most critical metrics for evaluating vaccine performance. An additional strength of the findings is the consistency of non-inferiority across diverse age groups. Young children are known to develop less durable immune responses following OCV administration and remain disproportionately vulnerable to severe cholera-related morbidity and mortality [23,24]. Nevertheless, subgroup analyses demonstrated stable immunogenic equivalence in both pediatric and adult populations, supporting the broad applicability of Euvichol formulations within preventive immunization programs and reactive outbreak campaigns.

The safety findings further reinforce the favorable tolerability profile of Euvichol formulations. No significant differences were identified in solicited adverse events or systemic symptoms between vaccine groups. The slight increase observed in unsolicited adverse events among Euvichol recipients was primarily attributable to mild upper respiratory tract infections deemed unrelated to vaccination by study investigators. Importantly, no vaccine-related serious adverse events, deaths, or life-threatening complications were identified across included trials. These findings are consistent with extensive postmarketing and clinical-trial evidence supporting the safety of killed whole-cell OCVs [25,26]. Maintaining this favorable safety profile remains essential for preserving public confidence and maximizing vaccine uptake during emergency outbreak responses.

## Limitations of Evidence

Several limitations inherent in the primary evidence should be acknowledged. First, the included studies primarily evaluated immunogenicity endpoints, including seroconversion and geometric mean titres (GMTs), rather than direct clinical efficacy outcomes such as culture-confirmed cholera infection. Although vibriocidal antibodies are accepted correlates of protection, they may not fully capture long-term mucosal immunity, indirect herd effects, or durability of protection in field settings.19 Second, one large multicenter trial employed an open-label design because differences in vaccine packaging prevented complete blinding [11]. This may have introduced reporting bias for subjective reactogenicity outcomes. Third, all included trials were conducted in Asian populations, including Bangladesh, India, Nepal, and the Philippines. Caution is therefore warranted when extrapolating these findings to African settings, where host immunity, nutritional status, environmental exposure, and cholera epidemiology may differ substantially. Additionally, vulnerable populations such as pregnant women, lactating mothers, and immunocompromised individuals were underrepresented in the available evidence base.

## Limitations of Review Processes

This review also faced several methodological constraints. The Baik et al. 2015 trial could not be included in pooled continuous GMT analyses because age-stratified standard deviations or confidence intervals were not reported in the primary publication. Although the study contributed to seroconversion analyses, its exclusion from continuous pooling may have slightly reduced the precision of GMT estimates. Moderate statistical heterogeneity was observed in the continuous GMT analysis for the O1 Ogawa serotype, likely reflecting differences in baseline immunity, endemic exposure intensity, and naturally acquired antibody levels across geographically diverse populations. Additionally, by restricting inclusion to randomized controlled trials to maximize internal validity, this review did not incorporate observational effectiveness studies or real-world campaign data that may further inform vaccine performance during humanitarian emergencies and mass-vaccination programs. A critical methodological distinction in our synthesis is the role of the Russo et al. (2018) trial. While it provides essential evidence for the internal stability and manufacturing consistency of the Euvichol platform across production scale-ups, the true comparison against the Shanchol benchmark rests on the other four trials. By isolating this study in our analysis, we ensured that the evidence for therapeutic equivalence was not confounded by manufacturing-specific equivalence data, while still acknowledging the study’s value in confirming the platform’s overall stability

### Implications

For policy and practice, these findings provide strong evidence supporting the interchangeable deployment of Euvichol, Euvichol-Plus, and Euvichol-S within global OCV stockpile programs. Confirmation of non-inferiority for Euvichol-S is particularly important because its simplified composition and scalable manufacturing process may substantially improve vaccine supply sustainability during periods of high global demand. These findings therefore support ongoing WHO and Gavi efforts to expand access to OCVs and advance the “Ending Cholera: A Global Roadmap to 2030” strategy.

Future research should prioritize long-term durability studies and real-world effectiveness evaluations of Euvichol-S in both endemic and epidemic settings. Additional studies involving pregnant women, immunocompromised individuals, and African populations are also needed to improve generalizability and ensure equitable protection across vulnerable groups. Finally, continued epidemiologic surveillance remains necessary to monitor the persistence of the O139 serogroup and confirm the long-term appropriateness of O1-focused vaccine strategies.

## Conclusion

This systematic review and meta-analysis suggests that contemporary Euvichol oral cholera vaccine formulations demonstrate comparable short-term immunogenicity and safety profiles to the Shanchol benchmark across randomized controlled trials conducted in cholera-endemic settings. Seroconversion responses against *Vibrio cholerae* O1 Inaba and Ogawa serotypes were consistently similar between vaccine platforms, with no major differences observed in solicited or serious adverse events. These findings support the continued use of Euvichol formulations as important components of global oral cholera vaccine supply strategies, particularly in the context of ongoing vaccine shortages and expanding cholera outbreaks. However, the interpretation of these results should consider several limitations, including the relatively small number of available trials, reliance on immunogenicity surrogate endpoints, variation in endpoint assessment timing, and limited long-term effectiveness data. Further post-marketing surveillance and effectiveness studies in diverse outbreak settings remain warranted.

## Contributors

Khalid Mohammed Al-Dhayani and Zaid Ahmad Hasan conceptualised the study and designed the search strategy. Khalid Mohammed Al-Dhayani and Khaled Ben Alwaleed Hassan Alastah, Mohammed Sameer Al-Eryani, and Sadiq M. Altbali independently performed data extraction and risk of bias assessments. Khalid Mohammed Al-Dhayani conducted the statistical analyses in RevMan, wrote the draft of the manuscript and provided critical review and logistical support. All authors had full access to all the data in the study, contributed to the interpretation of the results, critically revised the manuscript for important intellectual content, and had final responsibility for the decision to submit for publication.

## Data sharing

The de-identified participant summary data, full data extraction sheets, and the RevMan analytical files used for this meta-analysis will be made available upon reasonable request to the corresponding author immediately following publication.

## Acknowledgments

The authors would like to express their gratitude to ReseMeds Academy for providing the technical framework, statistical consulting, and research coordination that facilitated this study.

## Declaration of Generative AI in Scientific Writing

During the preparation of this work, the authors used AI-assisted technologies solely to improve language readability and structural formatting. After using this tool, the authors reviewed and edited the content as needed and take full responsibility for the content of the published article.

## ICMJE Attestation

All authors attest they meet the ICMJE criteria for authorship.

## Supplementary

Supplementary Appendix for

*Therapeutic Equivalence, Immunogenicity, and Safety of Euvichol Oral Cholera Vaccine Formulations Compared with the Shanchol Benchmark: A Systematic Review and Meta-Analysis*

Table of Contents:

1. Full Database Search Strategy
2. Methodological Notes for Meta-Analysis
3. Supplementary Table S1: Detailed Seroconversion Data Extraction
4. Supplementary Table S2: Continuous Data – Geometric Mean Titres (GMT) Data Extraction
5. Supplementary Table S3: Detailed Safety Data Extraction

## **1.** Full Database Search Strategy

The precise search string utilized across PubMed, MEDLINE via Ovid, Embase, Web of Science, and Scopus was:

> (Cholera OR "Vibrio cholerae") AND (Vaccine OR Vaccination OR "Oral Cholera Vaccine" OR OCV OR "Euvichol-Plus" OR "Euvichol-S" OR Shanchol OR Dukoral) AND ("Single-dose" OR "One-dose" OR "Two-dose") AND (Child OR Children OR Infant OR Toddler OR "Preschool Child" OR "1 year" OR "2 years" OR "3 years" OR "4 years" OR "5 years") AND (Immunogenicity OR Seroconversion OR "Antibody Titers" OR Effectiveness OR Efficacy OR Incidence OR Diarrhea OR Dehydration OR "Adverse Events").

## **2.** Methodological Notes for Meta-Analysis (Forest Plots)

To ensure statistical transparency and clinical homogeneity, the following data handling protocols were applied during the RevMan meta-analysis:

• **Design Heterogeneity (Russo et al. 2018):** One included study utilised an equivalence design comparing manufacturing variations (600L thimerosal-free vs. 100L fermenter) rather than a non-inferiority comparison against Shanchol. This study was isolated in a separate subgroup in all forest plots to assess internal manufacturing consistency.

• **Endpoint Timing (Chowdhury et al. 2022):** The primary immunogenicity endpoint for Chowdhury et al. 2022 was assessed at 7 days post-dose 2 (Day 21), whereas all other trials assessed responses at 14 days post-dose 2 (Day 28).

• **Data Aggregation:** For studies providing multiple paediatric strata (e.g., 1–5y and 6–17y), values were aggregated to represent a single 1–17y cohort for consistent cross-study comparison where required.

• **Derived Counts (Shah et al. 2023):** Event counts for Shah et al. 2023 were calculated by applying the reported percentage seroconversion rates to the per-protocol population size (N=103/arm) to allow inclusion in the Mantel-Haenszel dichotomous analysis.

• **Log-Transformation (GMTs):** Vibriocidal antibody titres follow a log-normal distribution; therefore, all GMT values and associated 95% CIs were natural log-transformed (ln) prior to analysis. The standard deviations for continuous data were derived from 95% CIs using the Cochrane formula: *SD*_log_ = √*N* × [ln(*UL*) − ln(*LL*)]/3.92

• **Missing Data (Baik et al. 2015):** Due to the absence of reported 95% CIs by age strata in the primary source for Baik et al. (2015), standard deviations could not be derived. These data points were excluded from the continuous GMT meta-analysis to avoid statistical error, but were retained for the dichotomous seroconversion meta-analysis.

• The Russo et al. (2018) study contributes to the overall proof of Euvichol’s stability but was methodologically separated to prevent conceptual inconsistency in the primary Euvichol-vs-Shanchol meta-analysis

